# *Mugil cephalus* Low-Density Lipoprotein Receptors: protein family characterization and identification of vitellogenin receptors

**DOI:** 10.1101/2025.03.25.645096

**Authors:** Miccoli Andrea, Randazzo Basilio

## Abstract

Vitellogenin receptors (VtgRs) are pivotal to the successful reproductive event in oviparous species by mediating the uptake of vitellogenin (Vtg) into developing oocytes, ensuring proper yolk formation and embryonic development. In this study, we characterized the low-density lipoprotein receptor (LDLR) family in flathead mullet *Mugil cephalus*, to which VtgRs belong, and placed a focus on identifying and analyzing vitellogenin receptors. Using a comprehensive approach that combined LDLR orthology inference across 13 species, protein domain analysis and 3D structure prediction, synteny evaluation, functional validation through RNA-seq-derived transcript models and phylogenetic analyses, we identified 87 LDLR family members and characterized two putative vitellogenin receptors belonging to the Lr8/VLDLR and Lrp13/LRX+1 subfamilies. These receptors exhibited conserved domain architectures, and the syntenic arrangements of the genes shared with orthologs in other teleost species as well as the ovary-specific expression profiles were consistent with the functional roles of VtgRs in mediating vitellogenin uptake during oocyte development. Phylogenetic analyses confirmed their evolutionary conservation, while structural modeling revealed key features related to ligand binding and functional specialization. The characterization of vitellogenin receptors in the flathead mullet not only advances the understanding of the reproductive biology in an aquaculture-relevant species but also lays the groundwork for future biotechnological interventions aimed at enhancing reproductive management and genetic improvement. Continued exploration of these avenues promises to improve the sustainability and economic viability of aquaculture practices, making it a fertile area for both basic research and applied biotechnology.

## Introduction

The low-density lipoprotein receptor (LDLR) family is a group of evolutionarily ancient cell surface receptors that is pivotal to lipid metabolism, cellular signaling, and development. These receptors are primarily known for their ability to bind and internalize lipoproteins, but they also participate in diverse biological processes, including nutrient transport, hormone uptake, and modulation of cellular signaling pathways. The LDLR family includes well-characterized members such as the LDL receptor (LDLR), LDL receptor-related proteins (LRPs), and very low-density lipoprotein receptor (VLDLR), among others. These share structural features, including ligand-binding domains, epidermal growth factor (EGF)-like repeats, and cytoplasmic tails containing NPxY motifs, which are involved in endocytosis and signal transduction ^1,2^, but distant members of the family display unrelated domain architectures encompassing, for instance, the vacuolar protein sorting-10 domain (VPS10) associated with Lr11, the CUB (for complement C1s/C1r, sea urchin epidermal growth factor and bone morphogenic protein-1) domain associated with the Lrp3, Lrp10, and Lrp12, or the MANEC domain associated with Lrp11 ^3^.

A major function of specific LDLR family members in the context of oviparous species’ reproductive physiology is the internalization of vitellogenin (Vtg), a yolk precursor protein essential for oocyte development and embryonic sustainment ^4,5^. Vtg is synthesized in the liver, secreted into the bloodstream, and for most forms ^6^, internalized by oocytes via vitellogenin receptor(s)-mediated endocytosis in clathrin-coated pits. Once internalized, Vtg is sequestered by growing oocytes and proteolytically cleaved into three smaller subunits, namely lipovitellin, phosvitin and β-component, which are stored in yolk globules or remain in the cytoplasm soluble fraction ^5,7,8^. The vitellogenin receptors, also known as VLDLR, are structurally and functionally homologous to other LDLR family members, featuring conserved ligand-binding and internalization domains that facilitate efficient endocytosis of yolk precursors. Beyond their role in Vtg uptake, LDLR family members, and in some cases directly VLDLR, are fundamental to key signaling pathways such as tyrosine kinases, Wnt, TGF-β, BMP and Reelin, essential for embryonic development and tissue homeostasis ^9^.

The flathead mullet (*Mugil cephalus*, Linnaeus, 1758) is an euryhaline teleost species, worldwide distributed in temperate and tropical shallow waters. In recent years it has gained relevant attention due to its significant potential in the aquaculture industry, driven by a number of factors, such as i) its fast growth and reduced reliance upon fishmeal and fish oil as feed ingredients, ii) the adaptability to a wide range of environmental conditions even in a global climate change scenario, iii) the need for diversifying the production in the Mediterranean region (https://cordis.europa.eu/article/id/169857-fishing-for-new-ways-to-expand-the-eus-aquaculture-industry) and iv) for ensuring access to high quality animal protein in developing countries ^10,11^. However, the cultivation of the species still mainly relies on the supply of wild-caught juveniles ^12,13^ since captivity breeding is limited by reproductive dysfunctions affecting early vitellogenesis, final oocyte maturation or spontaneous spawning depending on geographical location ^10,13–17^. Failing in gametogenesis in females has been attributed to impairment in vitellogenin uptake, thus resulting in arrested oocytes development ^18,19^, and hormone therapies based on gonadotropin releasing hormones and luteinizing hormone are used for oocyte maturation and spawning induction in wild-caught *M. cephalus* breeders ^17,19–24^. Reproductive dysfunctions in teleosts held in captivity were related to alterations in the endocrine control in the brain-pituitary-gonadal axis; however, a receptor-mediated interference in oocyte Vtg uptake cannot be ruled out.

To date, three native Vtg subtypes and seven Vtg-derived yolk proteins have been identified by column chromatography, electrophoresis and N-terminal peptide sequencing in *M. cephalus* vitellogenic ovaries and ovulated eggs protein extract ^25–28^. On the contrary, to the best of our knowledge, no studies on the species’ VtgRs are available so far.

In this paper, working exclusively *in silico* and employing an integrative approach that combines orthology inference across 13 species, homology searches, 3D protein structure modeling and domain analysis, syntenic evaluation, and transcriptional and phylogenetic reconstruction, we provide characterization of LDLR family members and particularly focus on putative vitellogenin receptors belonging to the Lr8/VLDLR and Lrp13/LRX+1 subfamilies.

## Methods

### Genomic resources

The proteomes of 13 species (*Mugil cephalus, Anguilla anguilla, Clupea harengus, Dicentrarchus labrax, Drosophila melanogaster, Danio rerio, Gadus morhua, Ictalurus punctatus, Oreochromis aureus, Oryzias sinensis, Seriola dumerili, Salmo salar* and *Salmo trutta*) belonging to 8 superorders (Acanthopterygii, Atherinomorphae, Clupeomorpha, Elopomorpha, Ostariophysi, Panorpoidea, Paracanthopterygii and Protacanthopterygii) and two classes (Actinopterygii and Insecta) were retrieved from Ensembl (release 113) or Ensembl Rapid (database version 110.1), with assemblies and accession numbers available in Table 1. Resources of *M. cephalus* were employed for LDLR data mining and characterization, vitellogenin receptor identification, protein structure modelling, synteny evaluation and expression and phylogenetic analysis, while the proteomes of the additional 12 species were employed for phylogenetic reconstruction.

**Table 1.** Assembly and accession number of genomic resources employed in this study. Species abbreviation, common name and scientific name, along with two taxonomic ranks are included.

| Abbreviation | Common name | Scientific name | Superorder | Order | Ensembl assembly | Accession |
| --- | --- | --- | --- | --- | --- | --- |
| mcep | Flathead mullet | *Mugil cephalus* | Acanthopterygii | Mugiliformes | CIBA_Mcephalus_1.1 | GCA_022458985.1 |
| aang | European eel | *Anguilla anguilla* | Elopomorpha | Anguilliformes | fAngAng1.pri | GCA_013347855.1 |
| char | Atlantic herring | *Clupea harengus* | Clupeomorpha | Clupeiformes | Ch_v2.0.2v2 | GCA_900700415.2 |
| dlab | European seabass | *Dicentrarchus labrax* | Acanthopterygii | Perciformes | dlabrax2021 | GCA_905237075.1 |
| dmel | Fruit fly | *Drosophila melanogaster* | Panorpoidea | Diptera | BDGP6.46 | GCA_000001215.4 |
| drer | Zebrafish | *Danio rerio* | Ostariophysi | Cypriniformes | GRCz11 | GCA_000002035.4 |
| gmor | Atlantic cod | *Gadus morhua* | Paracanthopterygii | Gadiformes | gadMor3.0 | GCA_902167405.1 |
| ipun | Channel catfish | *Ictalurus punctatus* | Ostariophysi | Siluriformes | ASM400665v3 | GCA_004006655.3 |
| oaur | Blue tilapia | *Oreochromis aureus* | Acanthopterygii | Perciformes | ZZ_aureus | GCA_013358895.1 |
| osin | Chinese medaka | *Oryzias sinensis* | Atherinomorphae | Beloniformes | ASM858656v1 | GCA_008586565.1 |
| sdum | Greater amberjack | *Seriola dumerili* | Acanthopterygii | Perciformes | Sdu_1.0 | GCA_002260705.1 |
| ssal | Atlantic salmon | *Salmo salar* | Protacanthopterygii | Salmoniformes | Ssal_v3.1 | GCA_905237065.2 |
| stru | Brown trout | *Salmo trutta* | Protacanthopterygii | Salmoniformes | fSalTru1.1 | GCA_901001165.1 |

### Orthology inference

Orthology was inferred in the *M. cephalus* proteome using the KofamScan software (https://github.com/takaram/kofam_scan), a gene function annotation tool based on the KEGG Orthology database (KO). K numbers were assigned by HMMER searches against a customized database of HMM profiles (https://www.genome.jp/ftp/db/kofam/, last update 2025-01-30), filtered with the “low density lipoprotein receptor” query, to sequences scoring above the predefined KO-dependent scoring criteria. The customized list of profiles against which the orthology search was performed, along with the corresponding threshold, F-measure and number of sequences from which each profile was built, is available in Table 2.

**Table 2.** Customized list of KEGG Orthology HMM profiles against which the orthology search was performed by HMMER, along with corresponding threshold score above which high confidence annotation was determined, F-measure and number of sequences from which the profile was built. For methodological details on score type, profile type and F-measure, the reader is redirected to 10.1093/bioinformatics/btz859.

| KO | symbol | name | threshold | score_type | profile_type | F-measure | nseq |
| --- | --- | --- | --- | --- | --- | --- | --- |
| K03068 | LRP5_6 | low density lipoprotein receptor-related protein 5/6 | 1129.27 | domain | all | 0.960461 | 1168 |
| K04550 | LRP1, CD91 | low-density lipoprotein receptor-related protein 1 (alpha-2-macroglobulin receptor) | 5253.87 | domain | all | 0.875475 | 730 |
| K08763 | OLR1 | oxidised low-density lipoprotein receptor 1 | 272.7 | domain | all | 0.895213 | 178 |
| K12473 | LDLR | low-density lipoprotein receptor | 1259.1 | domain | trim | 0.956793 | 479 |
| K12474 | LDLRAP1, ARH | low density lipoprotein receptor adapter protein 1 | 138.63 | full | all | 0.967019 | 1212 |
| K20049 | LRP1B | low-density lipoprotein receptor-related protein 1B | 7652.37 | full | all | 0.940485 | 470 |
| K20050 | LRP3_10_12 | low-density lipoprotein receptor-related protein 3/10/12 | 161.97 | domain | all | 0.994261 | 1263 |
| K20051 | LRP4, MEGF7 | low-density lipoprotein receptor-related protein 4 | 1210.23 | domain | all | 0.819646 | 881 |
| K20052 | LRP8, APOER2 | low-density lipoprotein receptor-related protein 8 | 1220.17 | domain | all | 0.828653 | 676 |
| K20053 | VLDLR | very low-density lipoprotein receptor | 1191.37 | domain | all | 0.801269 | 850 |

### Domain architecture and protein structure characterization

SMART ^29^ was employed for characterizing domain architectures of *M. cephalus* proteins assigned KO numbers with the greatest confidence (i.e. KofamScan results sorted alphabetically by KO and by score, and first result extracted). The theoretical isoelectric point (pI) and the predicted molecular weight (MW) of putative vitellogenin receptors were computed with the Expasy Compute pI/Mw ^30^. Protein structure of the putative vitellogenin receptors was predicted with atomic accuracy with the AlphaFold2 method ^31^ implemented in the COACH-D algorithm ^32^, which also predicted protein-ligand binding sites. The 3D protein structure was visualized and annotated with iCn3D ^33^.

### Vitellogenin receptor identification and sequence analysis

In-depth mining of the *M. cephalus* proteome against a NCBI-built non-redundant protein sequence database containing 501 Very Low Density Lipoprotein Receptor and 243 Lrp13 (also known as LRX+1) teleost proteins (Table S1a-b) was performed by blastp, BLAST+ suite v. 2.9 ^34^, setting the *e-value* argument to 1e-10 and the *max-target-seqs* argument to 1. The two putative vitellogenin receptors belonging to the Lr8 type (ENSMGCP00000018116.1 and ENSMGCP00000018117.1) were aligned with the slow/accurate pairwise alignment ClustalW algorithm (https://www.genome.jp/tools-bin/clustalw), setting gap open penalty to 10, gap extension penalty to 0.1 and using BLOSUM (Henikoff) as weight matrix.

### Synteny and RNA-seq reads-based validation of *M. cephalus* vitellogenin receptors

Functional validation of the putative vitellogenin receptors in *M. cephalus* was achieved by the evaluation of the syntenic arrangements of the encoding genes and by analyzing RNA-seq data obtained from spleen, kidney, gill, fertilized egg, male brain, male gonad, female brain and female gonad tissues (https://ftp.ensembl.org/pub/rapid-release/species/Mugil_cephalus/GCA_022458985.1/ensembl/rnaseq/). The information provided by the RNA-seq genebuild was exploited to identify the expression of tissue-specific transcripts.

### Phylogenetic analysis

The KofamScan pipeline run for *M. cephalus*, consisting in the orthology inference and the processing of the results (scripts available upon requests), was applied to the 12 additional species listed in Table 1. Proteins assigned K numbers with the highest confidence per KO per each species were fetched from the corresponding proteomes and submitted to a multiple sequence analysis using the *Super5* algorithm of the MUSCLE software v. 5.1 ^35^, which is designed to scale the Parallel Perturbed Probcons (PPP) algorithm by introducing divide-and-conquer heuristics, hence making it particularly suited for alignment of large datasets. The best-fit model of evolution of the protein alignment was estimated with ModelTest-NG ^36,37^. A phylogenetic tree with the maximum-likelihood (ML) optimality criterion and 1000 Felsenstein bootstrap replications was inferred with RAxML-NG ^38^ using 10 random and 10 parsimony-based starting trees. The computational resources necessary for estimating the best evolutionary model and the phylogenetic tree were provided free of charge by the CIPRES Science Gateway platform ^39^. iTOL v.7 ^40^ was used for tree visualization and annotation.

## Results and discussion

### *M. cephalus* proteome completeness and assessment

The *M. cephalus* proteome consists of 23352 proteins.

A measure of the *M. cephalus* proteome completeness was based on presence/absence of conserved genes in the Percomorphaceae lineage, a proxy for the ancestral gene repertoire in this clade, over 16080 conserved Hierarchical Orthologous Groups (HOGs). Representatives of these groups are expected to be present in the *M. cephalus* repertoire, and the proportion of missing HOGs proxied the proportion of missing genes in the total gene repertoire of the target proteome. Results on conserved single, duplicated, duplicated unexpected, duplicated expected and missing HOGs accounted for 15454 (96.11%), 306 (1.90%), 243 (1.51%), 63 (0.39%) and 320 (1.99%), respectively.

Whole proteome assessment was achieved by calculating the proportion of annotated protein-coding genes in the *M. cephalus* proteome that likely correspond to an actual protein-coding gene by comparing to the known gene family of the Percomorphaceae ancestral lineage. Genes in the “Consistent” category correspond to a gene family known to exist in the selected lineage; “Partial hit” proteins share similarity with proteins in known gene families on only part of their sequence, possibly indicating poorly defined gene models, structurally divergent genes or erroneous annotation; “Fragmented” proteins are those whose length is smaller than the proteins from the gene families they share similarity with (<50% median length), indicating likely fragmented sequences or erroneous annotations. Results of the whole proteome assessment accounted for 22975 (98.39%) consistent genes, 2444 (10.47%) partial hits and 353 (1.51%) fragmented.

Overall, these statistics define the completeness and reliability of all downstream analyses conducted.

### The LDLR family of *M. cephalus*

The orthology inference conducted by HMMER searches against a custom database of HMM profiles revealed that the proteome of the flathead mullet *M. cephalus* encompasses 87 low-density lipoprotein receptors (LDLR) sequences across 9 family member types. The most abundant one was the low-density lipoprotein receptor-related protein 8 (also known as Lrp8 or APOER2, KEGG Ortholog K20052, including 18 sequences), while the least abundant was the low-density lipoprotein receptor-related protein 3/10/12 (Lrp3_10_12, K20050, including 3 sequences) (Table S2). A representative depiction of the protein domain architecture of each of the LDLR family members is shown in Fig. 1, drawn from the sequences annotated with the highest confidence (i.e. highest score above the HMM threshold).

**Fig. 1.**
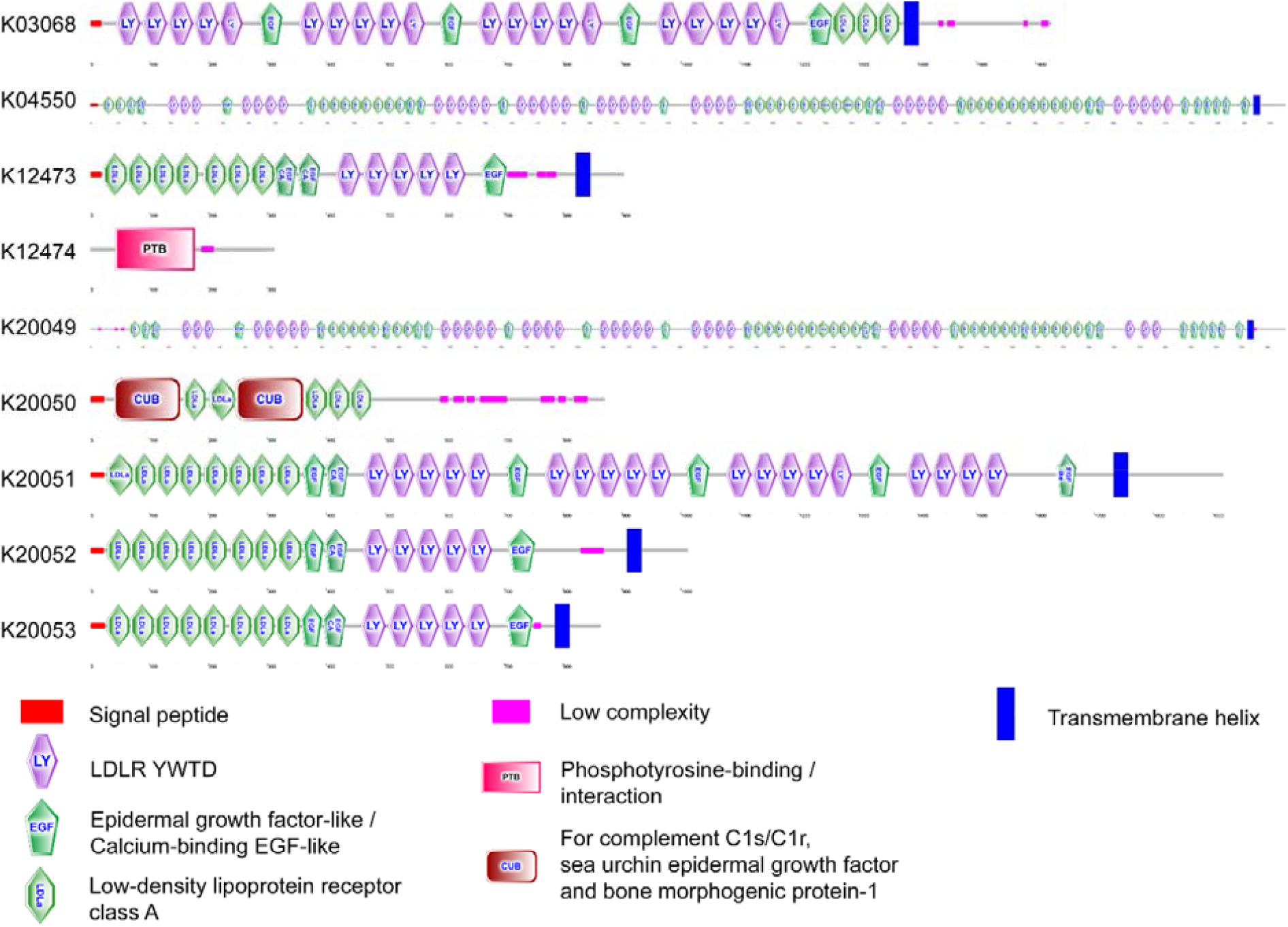
A representative depiction of the LDLR protein domain architecture, drawn from the protein sequences annotated with the highest confidence (i.e. highest score above threshold) per each class. K03068: ENSMGCP00000049234.1; K04550: ENSMGCP00000069134.1; K12473: ENSMGCP00000017067.1; K12474: ENSMGCP00000004882.1; K20049: ENSMGCP00000010065.1; K20050: ENSMGCP00000016101.1; K20051: ENSMGCP00000025192.1; K20052: ENSMGCP00000022056.1; K20053: ENSMGCP00000018116.1. LY: SMART accession number SM00135; EGF: SM00181; EGF_CA: SM00179; LDLa: SM00192; PTB: SM00462; CUB: SM00042. All LDLR types, except K04550 and K20049, are depicted in scale.

### *M. cephalus* putative vitellogenin receptors

Blastp searches of the *M. cephalus* proteome against a database specifically built with Lr8/VLDLR and Lrp13/LRX+1 teleost sequences yielded extremely significant results over three proteins, namely ENSMGCP00000018116.1, ENSMGCP00000018117.1 and ENSMGCP00000036233.1 (Table 3). Five hundred fifty-four out of 562 (98.4%) highly significant blastp hits were returned for ENSMGCP00000018116.1 and ENSMGCP00000018117.1 combined against Lr8/VLDLR, while 239 out of 321 (74.45%) hits were found for ENSMGCP00000036233.1 against Lrp13/LRX+1.

**Table 3.** Best *blastp* results of the homology search for the identification of vitellogenin receptors against a NCBI custom-built database of 501 Very Low Density Lipoprotein Receptor and 243 Lrp13 teleost proteins. Query: *M. cephalus* protein Ensembl ID; Subject: NCBI accession ID and species name of the first hit sorted by highest alignment score, i.e. score calculated from the sum of the rewards for matched amino acids and penalties for mismatches and gaps; Identity %: percent identity of the alignment.

| Query | Subject | Identity % | Encoding gene | Encoding transcript | AA length |
| --- | --- | --- | --- | --- | --- |
| ENSMGCP00000018117.1 | XP_070844659.1 [Chaetodon trifascialis] | 95.142 | *vldr* (ENSMGCG00000004079.1) | ENSMGCT00000018447.1 | 844 |
| ENSMGCP00000018116.1 | XP_070844658.1 [Chaetodon trifascialis] | 94.792 | *vldr* (ENSMGCG00000004079.1) | ENSMGCT00000018446.1 | 864 |
| ENSMGCP00000036233.1 | ASS77301.1 [Larimichthys crocea] | 80.017 | *si:dkey-88l16.3* (ENSMGCG00000012602.1) | ENSMGCT00000036864.1 | 1358 |

The architectures of ENSMGCP00000018116.1 and ENSMGCP00000018117.1 are composed of i) 8 low density lipoprotein class A receptor repeats containing the negatively-charged DxSDE conserved sequence (SMART accession number SM00192) for ligand-binding, a feature typical of other vertebrate vitellogenin receptors, ii) 3 epidermal growth factor like/calcium-binding EGF-like (SM00179 and SM00181); and iii) 5 YWxD repeats (SM00135) that form the β-propeller (Fig. 1, member K20053). ENSMGCP00000018116.1 has a theoretical isoelectric point (pI) of 4.76 and a predicted molecular weight (MW) of 94.74 kDa, while ENSMGCP00000018117.1 has a theoretical pI of 4.75 and a MW of 92.70 kDa, in line with molecular masses of Lr8-type vitellogenin receptors reported from other teleost species ^41^. The 2 Lr8 proteins differ only by the 20 AA-residue long peptide PDPSSKSPSKDDGKALIQPP (Fig. 2), encoded by the 3’ end of exon 16 and 57 out of 60 nt of exon 17 of transcript ENSMGCT00000018446.1: this peptide putatively corresponds to the O-linked sugar domain, as confirmed by the alignment against the VLDLR of *Homo sapiens* (GenBank accession # AAI36563.1) annotated by Babio and colleagues ^3^. The Lr8/VLDLR has been reported to exist in two isoforms: one containing an O-linked sugar domain and the other lacking it ^42^. The O-linked sugar domain supposedly blocks access to protease-sensitive sites, conferring protection against proteolysis, contributing to the stability of the receptor on the cell surface; instead, the variant lacking the O-linked sugar domain was confirmed to exhibit rapid cleavage from the cell surface, resulting in the release of a large amino-terminal VLDL receptor fragment in the extracellular space ^43^. Interestingly, no differences in ligand specificity or intracellular processing were observed between the two VLDLR variants, suggesting that the O-linked sugar domain primarily influences receptor stability and proteolytic susceptibility rather than functional interactions or trafficking ^44^. For these reasons, ENSMGCP00000018117.1 and ENSMGCP00000018116.1 can accordingly be referred to as *M. cephalus* Lr8-and Lr8+, respectively.

**Fig. 2.**
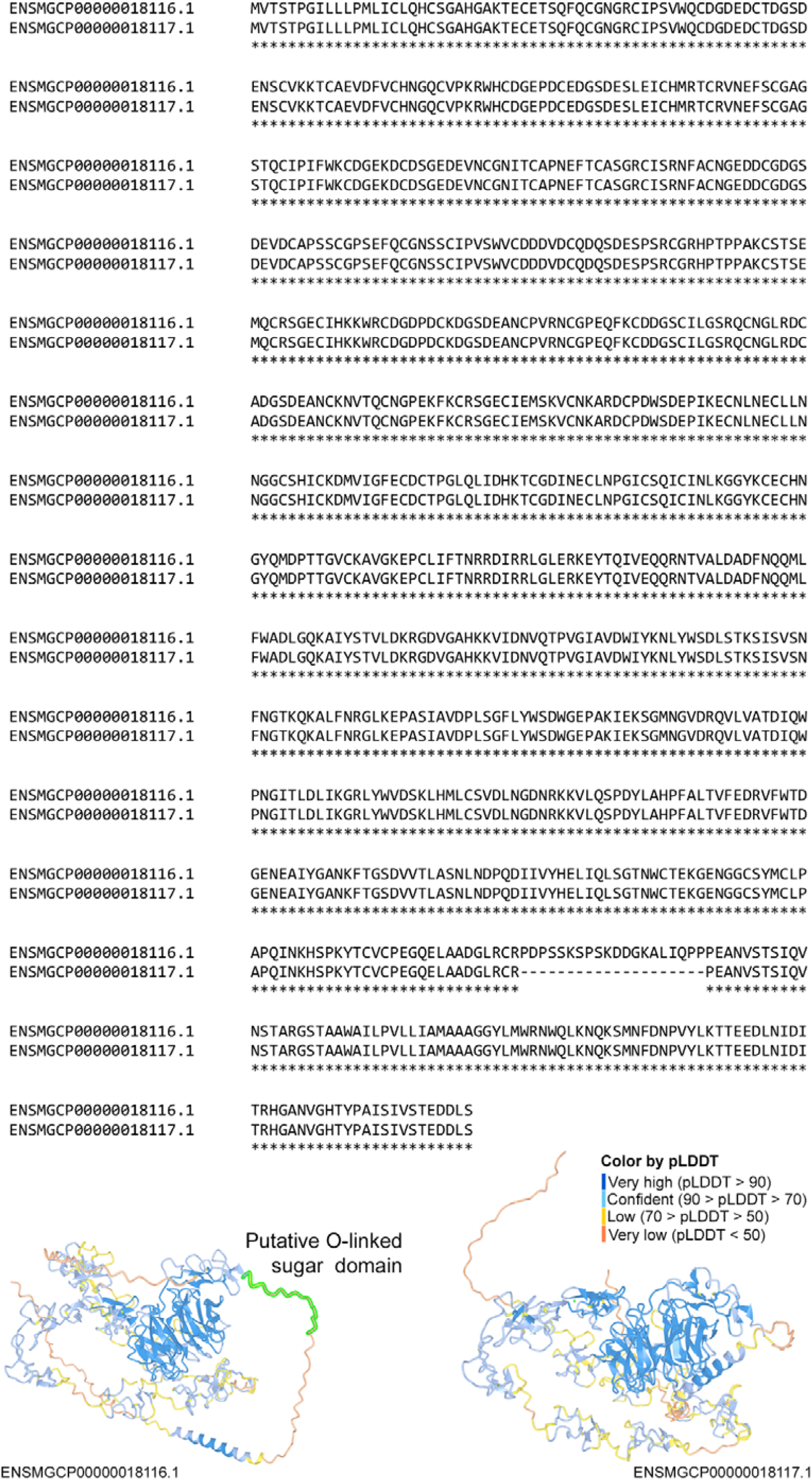
ClustalW alignment and 3D protein structure prediction of mcepLr8+ (ENSMGCP00000018116.1) and Lr8-(ENSMGCP00000018117.1), highlighting 100% residue conservation except for the 20-AA o-linked sugar domain region lacking in the Lr8-protein. Color annotation indicates per-residue measure of local confidence proxied by the predicted local distance difference test (pLDDT), scaled from 0 to 100, with higher scores indicating higher confidence and, accordingly, a more accurate prediction.

The architecture of ENSMGCP00000036233.1 displayed 3 additional LDL class A repeats compared to the above, of which one is characteristically located adjacent to the transmembrane domain. Such a feature is conserved in all Lrp13 orthologs despite the number of N-terminal LDLa domains ranges from 7 to 10 among teleost species ^45^. Also, the sequence presents 2 additional EGF/EGF CA domains preceding the C-terminal LDL repeat and the transmembrane domain, and 3 additional YWTD repeats (Fig. 3), in agreement with previous findings ^46^. ENSMGCP00000036233.1 has a theoretical pI of 4.79 and a MW of 147.39 kDa. It must be noted that a shift between the MW predicted from the polypeptide sequence and the apparent mass detected in the ovarian membrane appears to be a standard feature of Lrp13, as per observations in phylogenetically distant species such as cutthroat trout *Oncorhynchus clarki* and white perch *Morone americana* ^45,46^.

**Fig. 3.**
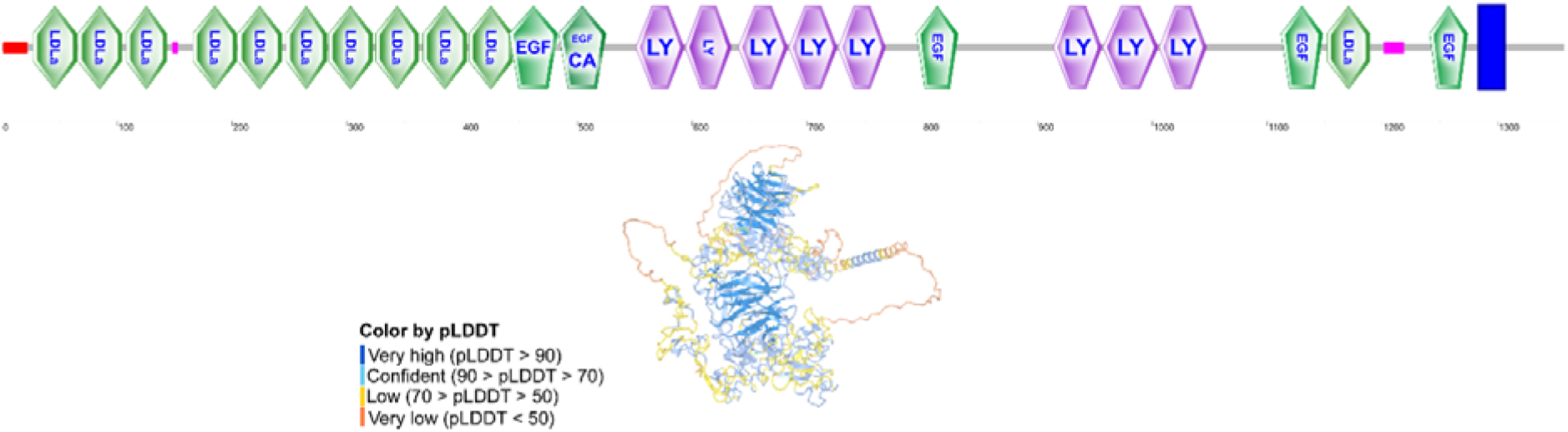
mcepLrp13 protein domain architecture and 3D protein structure prediction. For SMART accession numbers see the legend of Fig. 1. Protein model color annotation as per Fig. 2.

Lr8- and Lr8+ are encoded by two of the ten splice variants of gene ENSMGCG00000004079.1, namely ENSMGCT00000018447.1 and ENSMGCT00000018446.1, respectively. Lrp13 is instead translated from transcript ENSMGCT00000036864.1 (gene ENSMGCG00000012602.1). In addition to the protein architecture, proof-of-concept of their designation as vitellogenin receptors was provided by the evaluation of the syntenic arrangements of *vldlr/lr8* and *lrp13* loci as well as by transcriptional information. Both *vldlr/lr8* and *lrp13* (identified as si:dkey-88l16.3 in the CIBA_Mcephalus_1.1 assembly) genes share genomic synteny with orthologues in *A. anguilla, D. rerio, O. niloticus, O. latipes*, three-spined stickleback *Gasterosteus aculeatus* and spotted gar *Lepisosteus oculatus* ^3,45,47^. The flathead mullet *lr8* gene (chromosome 19, position 20188293-20232020) is found on the forward strand and is flanked by *sh3bp2* (SH3-domain-binding protein 2) and *kcnv2a* (potassium channel subfamily V member 2a), while *spp1* (secreted phosphoprotein 1) and *pum3* (pumilio RNA-binding family member 3) are the two closest loci on the reverse strand (Fig. S1); instead, the flathead mullet *lrp13* (chromosome 8, position 22833187-22843398) is on the reverse strand and is neighbored by *tia1* (cytotoxic granule associated RNA binding protein) and *upb1* (beta-ureidopropionase 1), while *rchy1* (ring finger and CHY zinc finger domain containing 1) and *smad2* (SMAD family member 2) are the two closest loci on the forward strand (Fig. S2). The syntheny of vitellogenin receptors across teleost species highlights their fundamental role in reproductive biology.

RNA-seq genebuild confirms Lr8- and Lrp13 to be the ovarian protein isoforms expressed in *M. cephalus*, as indicated by the female gonad RNA-seq gene model matching the exon structure of transcript ENSMGCT00000018447.1 and ENSMGCT00000036864.1, respectively (Fig. 4A-B). This visualization highlights the differences between the reference annotation and RNA-seq-derived transcript models, providing insights into tissue-specific gene expression and alternative splicing. Lr8 expression data from additional tissues is reported in Fig. S3: the Lr8-transcript was detected as the sole isoform in fertilized eggs, while both Lr8 splice variants were found in spleen, kidney and the brain of both male and female specimens. In brain, Lr8 expression may relate to signal transduction or general lipid metabolism in the nervous system ^41^, while transcriptional signatures in the fertilized egg (i.e. embryo) are likely due to vitellogenins having multiple roles in regulating buoyancy, development, nutrition and hatching ^48^. The Lr8 lacking the O-linked sugar domain was reported to be the dominant ovarian vitellogenin receptor isoform in a multitude of oviparous vertebrates, including teleosts ^41,47,49^, and evidence across a broad range of taxa report the ovarian form lacking the O-linked sugar domain to be responsible for vitellogenin uptake ^42,50,51^. On the other hand, Lr8+, whose expression may occur in the ovary as well ^3^, is hypothesized to act as a somatic VLDLR ^5,42,50^. Lrp13 is functionally translated solely in the ovary.

**Fig. 4.**
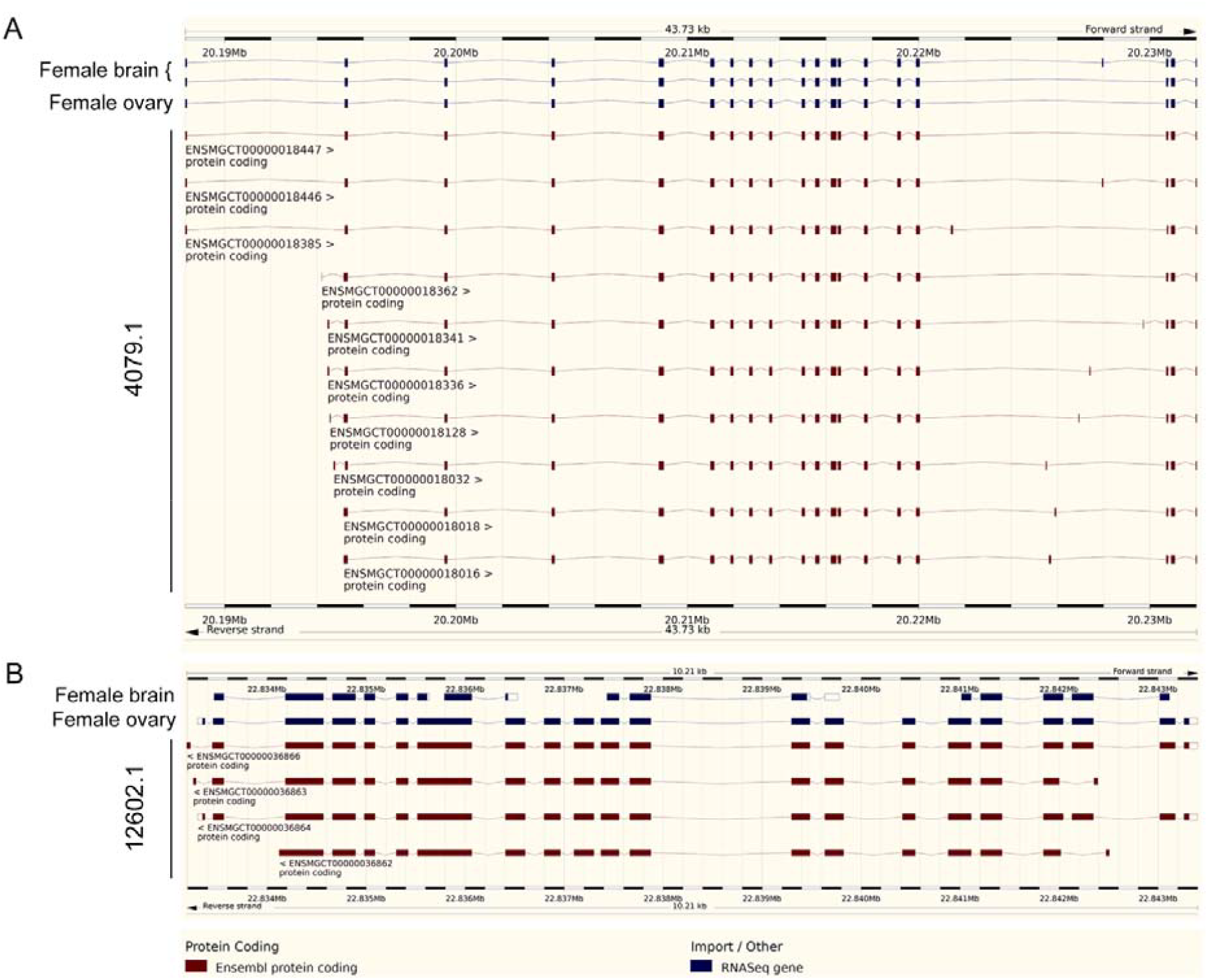
Visualization of gene models from the Ensembl Genome Browser. The Ensembl genome genebuild of genes ENSMGCG00000004079.1 (A) and ENSMGCG00000012602.1 (B) (red features) are overlaid with RNA-seq-based gene predictions obtained from female brain and female gonad transcriptomes (blue features). Gene models are displayed with exon-intron structures, where boxes represent exons and connecting lines represent introns. Solid blue-filled boxes are coding sequences, while empty boxes indicate non-coding untranslated regions (UTRs).

Historically, only the Lr8-type LDLR was identified and characterized as a vitellogenin receptor in oviparous vertebrates ^52^, even though multiple ovarian membrane proteins that specifically bind vitellogenin had been detected by ligand blotting in salmonid and perciform species ^53–58^ before additional VtgR types were formally characterized ^45,59^. The characterization of VtgRs in fish revealed functional versatility ^58^, which is likely an adaptive response to ensure efficient oocyte nourishment in different environmental conditions. Herein, RNA-seq supports the hypothesis that the ENSMGCT00000018447.1 (i.e. Lr8-) and ENSMGCT00000036864.1 (Lrp13) splice variants be the two actual functional flathead mullet vitellogenin receptors, supporting the oocyte uptake of different forms of vitellogenins that result in varying degrees of storage and/or proteolysis of corresponding yolk proteins along ovarian maturation and embryonic development. The proportions of yolk vitellogenin subtypes vary significantly among teleosts depending on their reproductive strategies and early life histories ^60,61^; in the flathead mullet, a pelagic spawner possessing three vitellogenin subtypes ^27^, the oocyte accumulation ratio of VtgAa, VtgAb and VtgC is 4:13.3:1 ^62^. Reporting Lrp13 in this species reinforces the hypothesis about the existence of a system for differential uptake that is likely receptor-mediated, with Lr8- and Lrp13 supposedly preferentially binding VtgAb and VtgAa ^63^, and VtgC possibly entering via common endocytosis or in association with other proteins ^6^.

### Phylogenetic analysis

The best-fit amino acid replacement model for the protein alignment according to Bayesian Information Criterion (BIC), Akaike Information Criterion (AIC) and its small-sample equivalent (AICc) was “JTT+I+G4”, with negative log-likelihood values of -145296.0767, -145296.0767 and - 145296.0767, respectively. The phylogenetic reconstruction in Fig. 5 illustrates the evolutionary relationships of LDLRs in several fish species belonging to 7 superorders. It was build with 4 insect LDLR sequences intended as outgroups, and is also annotated with the overall number of sequences annotated as one of the KO types in Table 2 for each of the proteomes considered.

**Fig. 5.**
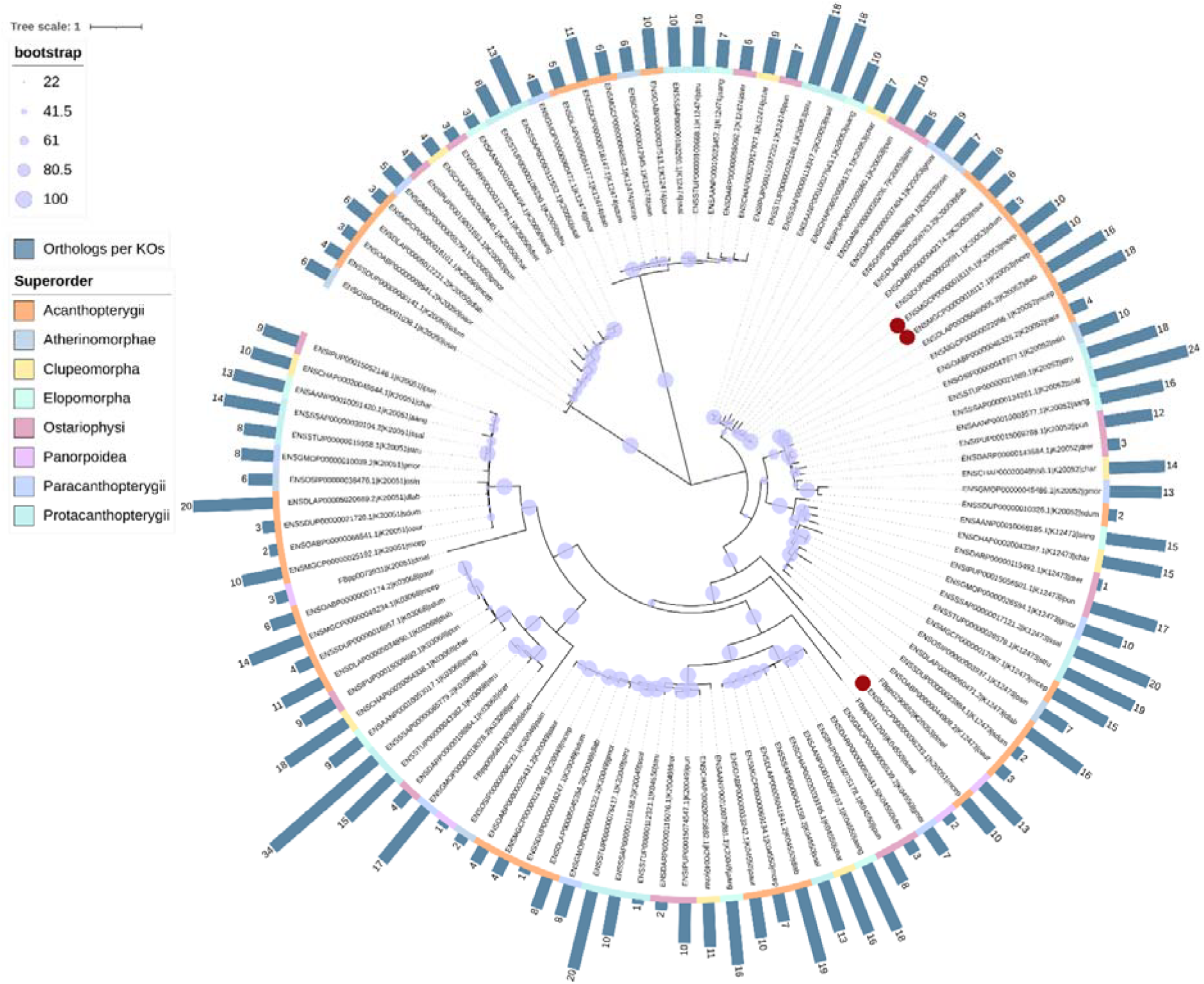
Phylogenetic reconstruction of the LDLR family. The Maximum Likelihood tree was constructed using the JTT+I+G4 model of evolution and 1000 bootstrap replicates. The bar chart indicates the overall number of sequences annotated as one of the KO types in Table 2 (inclusive of the sequence employed for MSA and phylogenetic tree construction), while the colored strip refers to the Superorder taxonomic lineage to which the investigated species belongs. The raw data used for the bar chart annotation is available in tabular formato per KO per species in Table S2. The three sequences herein characterized are highlighted by red circles.

The tree followed an expected pattern of divergence, where orthologs clustered according to KO types and known phylogenetic relationships as proxied by taxonomic lineages.

All *M. cephalus* LDLRs always clustered within clades that included other Acanthopterygii species, here represented by European sea bass *Dicentrarchus labrax*, greater amberjack *Seriola dumerili* and Blue tilapia *Oreochromis aureus*, confirming shared evolutionary history within the superorder and suggesting that LDL receptors retained common structural and functional characteristics.

Proteins displaying the same architecture, and therefore belonging to the same LDLR family member, across various superorders always clustered together, suggesting the retaining of functional specialization within each receptor type. Interestingly, the 4 sequences of *D. melanogaster* displaying the K20051, K03068, K04550 and K20053 architectures, were retained within the branches formed per KO type encompassing teleost LDLR proteins, despite in a distinct branch with high probability support (100 for K03068, 96 for K04550 and K20051, and 90 for K20053), demonstrating that fruit fly LDLR sequences of a given architecture are more similar to teleost sequences of the same architecture than they are to other fruit fly LDLR members. This emphasized the structural and functional conservation of LDL receptors across metazoans, with species estimated to have a phylogenetic separation of 10^9^ years of evolution harboring several strikingly similar family members in both structure and exon/intron organization ^64^, while highlighting the evolutionary distance between invertebrate and vertebrate receptor subtypes, also with regards to numerosity (i.e. number of members belonging to each of the LDLR types) and, presumably, multifunctional gain and increasing specialization allowing proper intercellular communication *sensu latu* in addition to lipid transport. The evolutionary history of the LDLR family, which gradually expanded along phylogenetic evolution ^65^, suggests that these receptors arose rapidly by repeated duplication and concatenation of exon/intron blocks from a single primordial precursor gene with the emergence of multicellular organisms, adapting to various physiological functions beyond lipid metabolism ^64^.

Sequences annotated as K12474 and K20050 displayed the greatest diversity between one another and with respect to all other LDLR types. This was especially evident in the unrooted visualization of the phylogenetic tree (data not shown), but is not surprising since their architecture, which encompasses PTB and CUB domains, differs markedly from all others. Proteins with such structures were regarded to as distant members of the LDLR family: proteins encoding phosphotyrosine binding domains function as adaptors or scaffolds to organize the signaling complexes involved in wide-ranging physiological processes including neural development, immunity, tissue homeostasis and cell growth ^66^, while proteins with the CUB module appear to be mainly extracellular and plasma membrane-associated proteins, many of which are developmentally regulated ^67^. Members of the LDLR family play a role in a diverse set of biological functions that transcend lipid metabolism, regulating for instance lipoprotein trafficking, synaptic plasticity, signal transduction, cell migration, and cellular growth regulation and cancer also in the developing and adult nervous system ^68^. Recently, several family members were found involved in amyloid precursor protein processing and β-amyloid secretion, demonstrating their participation in Alzheimer’s disease pathogenesis and neurodegeneration ^69^.

The probability support of the distinct allocation of sequences annotated as K20053 from all other KO types was 34, demonstrating that the evolutionary relationships of LDLR protein architectures presenting the LDLR YWTD domain, the Epidermal growth factor-like domain, the calcium-binding EGF-like domain and the low-density lipoprotein receptor class A domain cannot be resolved unequivocally. The importance of employing bioinformatic methods that implement probabilistic models such as the hidden Markov models for correctly annotating large protein families and detecting remote homologs as sensitively as possible is warranted.

The putative *M. cephalus* vitellogenin receptors herein characterized were included in the multiple sequence alignment for phylogenetic reconstruction, although only ENSMGCP00000018116.1 matched the criteria for inclusion in the dataset. In detail, protein ENSMGCP00000018117.1 was not originally fetched for phylogenetic reconstruction because a *M. cephalus* protein with the K20053 architecture was already included in the dataset, and neither was protein ENSMGCP00000036233.1 because low-confidently annotated as K20051 (“low-density lipoprotein receptor-related protein 4”, score of 648.1 over a threshold of 1210.23, e-value of 1.2e-193) by HMMER, and only later confirmed as Lrp13 by a blastp search. The 2 Lr8 variants clustered together with all other K20053 proteins, and Lrp13 was identified as outgroup (support probability of 92) to a branch including 47 additional leaves displaying 4 protein architectures. Importantly, the two *M. cephalus* vitellogenin receptor types clustered distantly from one another, suggesting that the earliest divergence within the LDLR family genes may have occurred between LRPs and LRs.

## Conclusions and future perspectives

This study provided a comprehensive characterization of the LDLR family in the flathead mullet, specifically focusing on the identification of its vitellogenin receptors. The approach integrated orthology inference across 13 species with the analysis of protein domain architectures, homology searches, evaluation of syntenic arrangements of the vitellogenin receptor-encoding genes and an RNA-seq-based analysis of tissue specificity, overall allowing the confident identification of VtgRs in the mullet.

Vitellogenin receptors mediate the uptake of vitellogenin into developing oocytes and eggs, a critical process for egg maturation and for the sustainment of the developing embryo: in this context, the molecular characterization of distinct vitellogenin receptors provides valuable information for deepening the knowledge on the reproductive physiology of a teleost species with huge aquaculture potential and improving broodstock management through optimized conditioning and synchronization of spawning. VtgRs also offers promising avenues for selective breeding and genetic improvement as the identification of favorable allelic variations associated with efficient vitellogenin uptake may serve as molecular markers in breeding programs, thereby enabling the selection of flathead mullet specimens with superior reproductive performance. Such advancements could significantly boost productivity and sustainability in aquaculture operations of this species.

The findings herein reported have important implications for adapting captive breeding strategies, as reproductive events in captivity often fail due to stress and altered environmental cues. Indeed, under captive conditions, flathead mullet breeders of both sexes suffer from reproductive dysfunctions. By deepening our understanding of the pathways mediated by VtgRs, future works may focus on developing approaches that better mimic natural reproductive cycles, thereby mitigating the challenges associated with captivity and elevating egg quality, fecundity and fertilization rates, even though direct, applied case studies specifically exploiting VtgRs in aquaculture are currently limited.

Additionally, the characterization of these receptors establishes a baseline for assessing the impact of environmental stressors such as pollutants and endocrine disruptors on reproductive health, opening to their use as biomarkers either alone or in conjunction with the more frequently-investigated vitellogenin in case the latter fails to provide unequivocal information ^70,71^.

## Acknowledgements

This work has been financially supported by the National Recovery and Resilience Plan (NRRP), Mission 4 Component 2 Investment 1.4 - Call for tender No. 3138 of 16 December 2021, rectified by Decree n.3175 of 18 December 2021 of Italian Ministry of University and Research funded by the European Union – NextGenerationEU (Award Number: Project code CN_00000033, Concession Decree No. 1034 of 17 June 2022 adopted by the Italian Ministry of University and Research, CUP D33C22000960007, Project title “National Biodiversity Future Center - NBFC”).

## CRediT authorship contribution statement

AM: Conceptualization, Methodology, Investigation, Visualization, Formal analysis, Data curation, Writing – original draft. BR: Visualization, Validation, Writing – editing and review.

## Data availability

All genomic resources analyzed in this study are publicly available and accession IDs were reported in Table 1. The scripts built for data manipulation and processing are available upon request to the corresponding author.

## Notes

### Competing Interest Statement

The authors have declared no competing interest.

