## Supplementary figures and images for "*Mugil cephalus* Low-Density Lipoprotein Receptors: protein family characterization and identification of vitellogenin receptors"

### Fig. S1

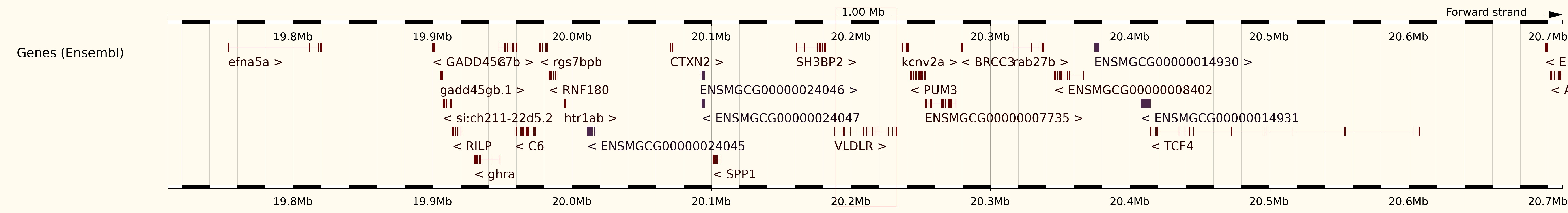

### Fig. S2

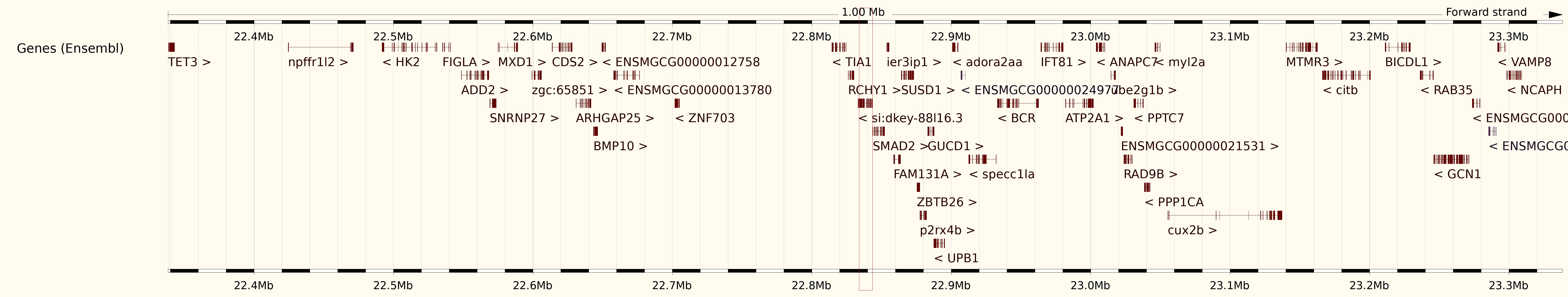

### Fig. S3

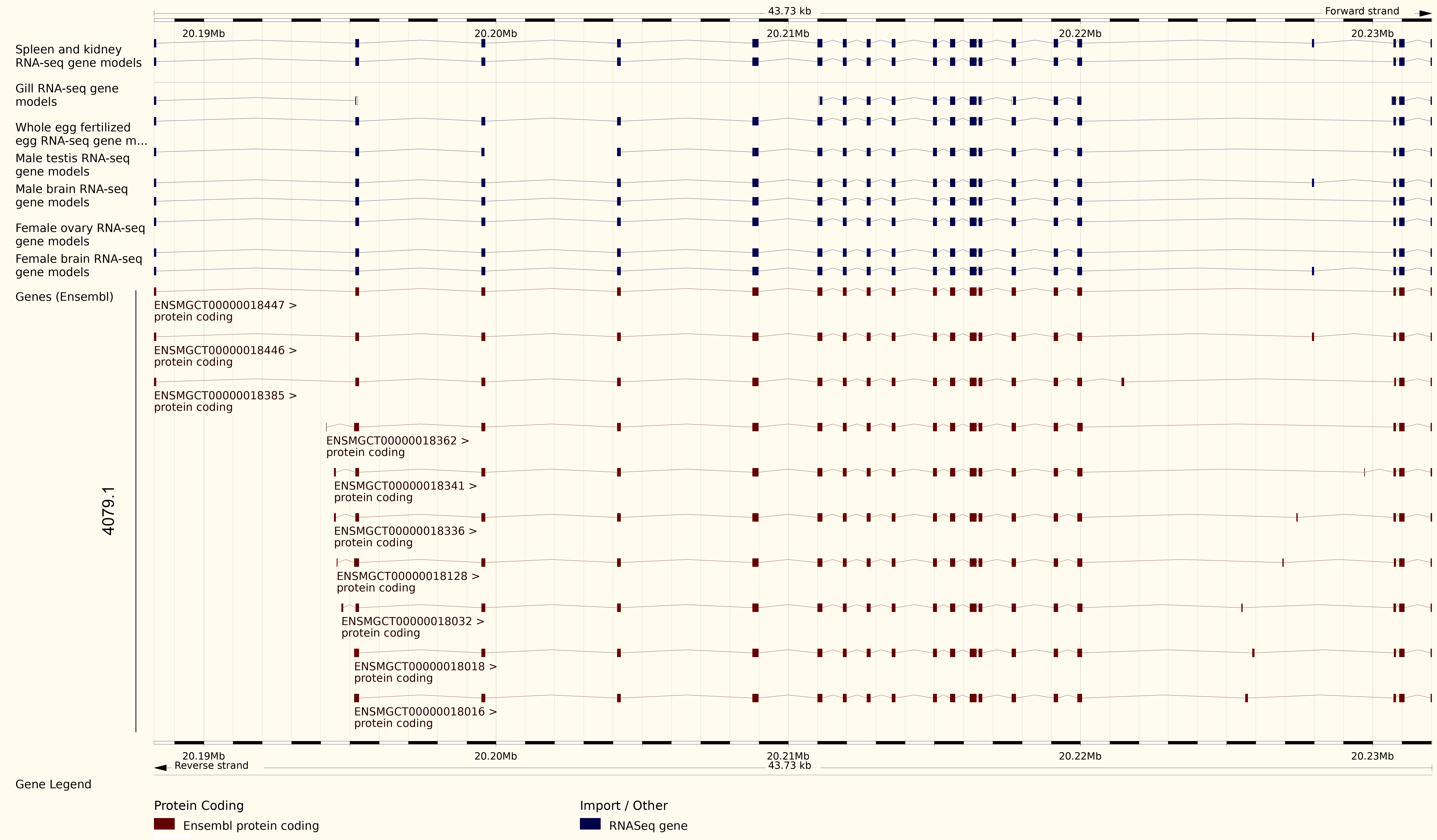
